# CD150-dependent hematopoietic stem cells sensing of *Brucella* instructs myeloid commitment

**DOI:** 10.1101/2021.03.31.437872

**Authors:** Hysenaj Lisiena, De Laval Bérengère, Arce-Gorvel Vilma, Bosilkovski Mile, Gonzalez Gabriela, Debroas Guillaume, Sieweke Michael, Sarrazin Sandrine, Gorvel Jean-Pierre

## Abstract

So far, hematopoietic stem cells (HSC) are considered the source of mature immune cells, the latter being the only ones capable of mounting an immune response. Recent evidence shows HSC can also directly sense cytokines released upon infection/inflammation and pathogen-associated molecular pattern interaction, while keeping a long-term memory of previous encountered signals. Direct sensing of danger signals by HSC induces early myeloid commitment, increases myeloid effector cell numbers and contributes to an efficient immune response. Here, using specific genetic tools on both host and pathogen sides, we show that HSC can directly sense B. abortus pathogenic bacteria within the bone marrow via the interaction of the cell surface protein CD150 with the bacterial outer membrane protein Omp25, inducing efficient functional commitment of HSC to the myeloid lineage. This is the first demonstration of a direct recognition of a live pathogen by HSC via CD150, which attests of a very early contribution of HSC to immune response.

**SUMMARY:** This work provides first evidence HSC directly sense Brucella abortus via the bacterial outer membrane protein Omp25 and the HSC surface receptor CD150, leading to functional commitment of HSC to myeloid lineage and very early initiation of immune response.

## INTRODUCTION

Few pathogens are capable of colonizing the bone marrow (BM) [1–5], the niche of haematopoietic stems cells (HSCs) responsible for initiating the production of myeloid progenitors and mature blood-forming cells [6]. *Brucella abortus*, (Gram-negative bacterium responsible for the re-emerging zoonosis of brucellosis) is able to persist for months in the BM [7]. Human brucellosis patients suffer from haematological abnormalities suggesting that Brucella in the BM may affect hematopoietic development [8]. HSCs can respond to an infection through pathogen-elicited cytokines or directly via pathogen recognition receptors (PRR) [9, 10]. CD150 is a key marker of long term HSC (LT-HSCs) [11] and a microbial sensor in macrophages [12] and dendritic cells [13]. Here, we show that HSCs within the BM directly sense the outer membrane protein Omp25 of live *Brucella* through the CD150 receptor. Our *in vivo* and *ex vivo* data demonstrate that *B. abortus* modulates hematopoiesis by transiently augmenting the production of myeloid cells via CD150. This is the first demonstration of a direct recognition of a pathogen by HSCs via CD150.

## RESULTS

To investigate the consequences of BM infection by a pathogen known to persist for extended periods of time in the hematopoietic niche, we used *Brucella abortus*. We previously demonstrated that this pathogenic bacterium persists for months in murine BM [7]. We began by evaluating if bacteria present in the BM at 8 days post-infection (acute phase of infection) and at 30 days post-infection (chronic phase of infection) are still virulent. For this, we transplanted BM cells of infected mice at those time point into recipient mice (Fig. 1a). Spleen (the natural reservoir for *B. abortus*) and BM cells were harvested eight weeks after transplantation, and bacterial colony forming units (CFU) were enumerated (Fig.1b). The equivalently high number of bacteria in spleen and bone marrow at 8 weeks post-transplantation shows that the BM hematopoietic environment is permissive for stable infection and replication of *Brucella*.

**Figure 1:**
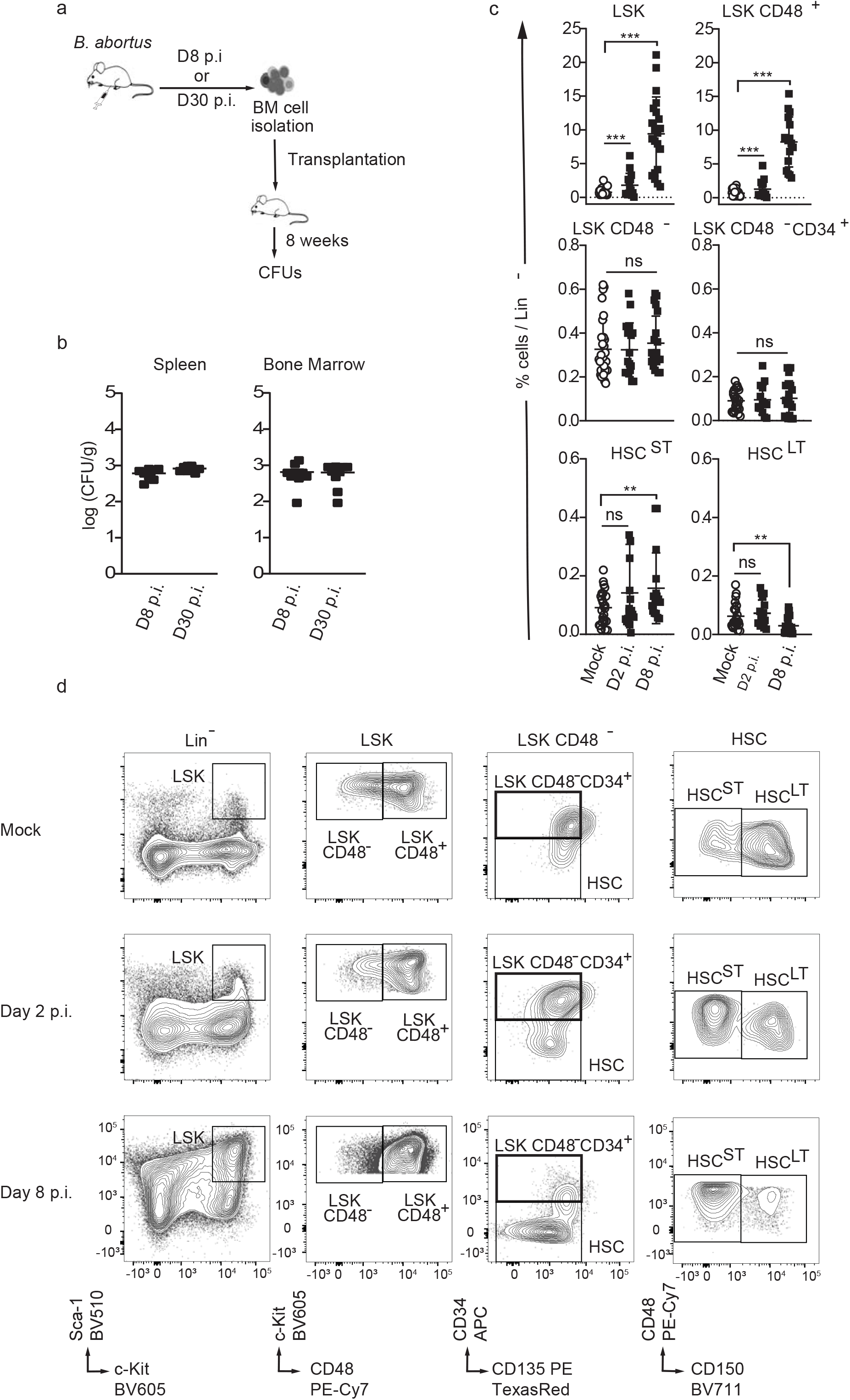
*Brucella abortus* persists in the BM and affects HSPC homeostasis. **a)** Experimental scheme: Mice were intraperitoneally inoculated with 1×10^6^ CFU of wild-type *B. abortus*. BM cells were isolated from femur and tibia of the infected mice, resuspended in PBS and transplanted into previously lethally irradiated mice. 8 weeks after transplantation CFU per gram of organ were enumerated from spleens and bone marrow (BM). **b)** Enumeration of CFUs per gram of spleen and BM at 8 weeks post transplantation (n=7). **c-d)** C57BL/6J wild-type (*wt*) mice were intraperitoneally inoculated with 1×10^6^ CFU of wild-type *B. abortus*. Two, eight and thirty days later, FACS analyses were performed for BM cells. Representative FACS profiles **(d)** and frequency of LSK (lin^-^,Sca^+^,cKit^+^) (from left to right, n=30; 15; 21), LSK CD48^+^ (lin^-^,Sca^+^,cKit^+^ CD48^+^) (from left to right, n=28; 14; 17), LSK CD48^-^ (lin^-^,Sca^+^,cKit^+^ CD48^-^) (from left to right, n=30; 17; 20), LSK CD48^-^ CD34^+^ (lin^-^,Sca^+^,cKit^+^ CD48^-^, CD34^+^, CD135^-^) (from left to right, n=24; 14; 19), HSC^ST^ (lin^-^,Sca^+^,cKit^+^ CD48^-^, CD135^-^ CD34^-^,CD150^-^) (from left to right, n=25; 17; 16), HSC^ST^ (lin^-^,Sca^+^,cKit^+^ CD48^-^, CD135^-^ CD34^-^,CD150^+^) (from left to right, n=22; 17; 22) in lineage negative fraction of BM **(c)** for PBS treated (Mock ○) and infected mice (■). Data were obtained from distinct samples and from 5 independent experiments, each with at least n=3 animals per condition, are shown and mean ± SEM is represented by horizontal bar. Significant differences from mock are shown. *** *P* < 0.001, ** *P*< 0.01, * *P*< 0.05. Absence of *P* value or ns, non-significant. Since data followed normal distribution, P-Value were generated using Brown-Forsyth followed by ANOVA Welch test.

To further investigate the consequences of BM infection by *B. abortus* on hematopoietic stem cell biology, we analysed the distribution of the hematopoietic stem and progenitor cell compartment (HSPC) by flow cytometry. Absolute numbers of total BM cells or lineage negative progenitors (Lin^-^) were not affected by *B. abortus* infection (Supplementary Fig. 1a, b). However, *Brucella* infection induced major phenotypic cell surface marker changes in the HSPC compartment (Fig. 1c, d and Supplementary Fig. 1c for gating strategy), similar to those observed after challenges with PAMPs or attenuated vaccines [10, 14–16]. HSPC (LSK: Lin^-^, Sca^+^, cKit^+^) expansion was observed at the onset of infection (Day 2 p.i., Fig. 1c, d) and was even more pronounced during the acute phase of infection (Day 8 p.i., Fig. 1c, d). LSK expansion was mainly due to the significant increase of CD48^+^ multipotent progenitors (LSK CD48^+^) and, to a lesser extent, the increase of short term HSC (ST-HSC: LSK, CD34^-^, CD135^-^, CD48^-^, CD150^-^). In addition, the long-term HSC population (LT-HSC: LSK, CD34^-^, CD135^-^, CD48^-^, CD150^+^) slightly decreased (Fig. 1c, d). Overall, these data suggest that the presence of *Brucella* in the BM perturbs HSPC homeostasis and leads to an increased output of early multipotent progenitors.

Infection and inflammation have been shown to release signals, such as cytokines, that are able to induce the differentiation of HSC towards the myeloid lineage as evidenced by early up-regulation of the myeloid master regulator PU.1 [17–20]. To further investigate whether *B. abortus* infection can induce an early commitment towards the myeloid lineage in HSC, we infected *Pu.1*^+/*GFP*^ reporter mice harbouring enhanced green fluorescent protein (GFP) knocked into the PU.1 locus [21, 22] and then analysed GFP expression in HSC (LSK, CD34^-^, CD135^-^, CD48^-^) at 2 days post-infection. GFP expression in HSC was used as a read-out of early HSC activation/commitment during *Brucella* infection (Fig. 2a). GFP was upregulated 30 to 40% in BM HSCs of *B. abortus-infected* WT mice (Fig. 2b, left panel), confirming an induction of PU.1 expression and consequently a change in HSC fate after *Brucella* infection.

**Figure 2:**
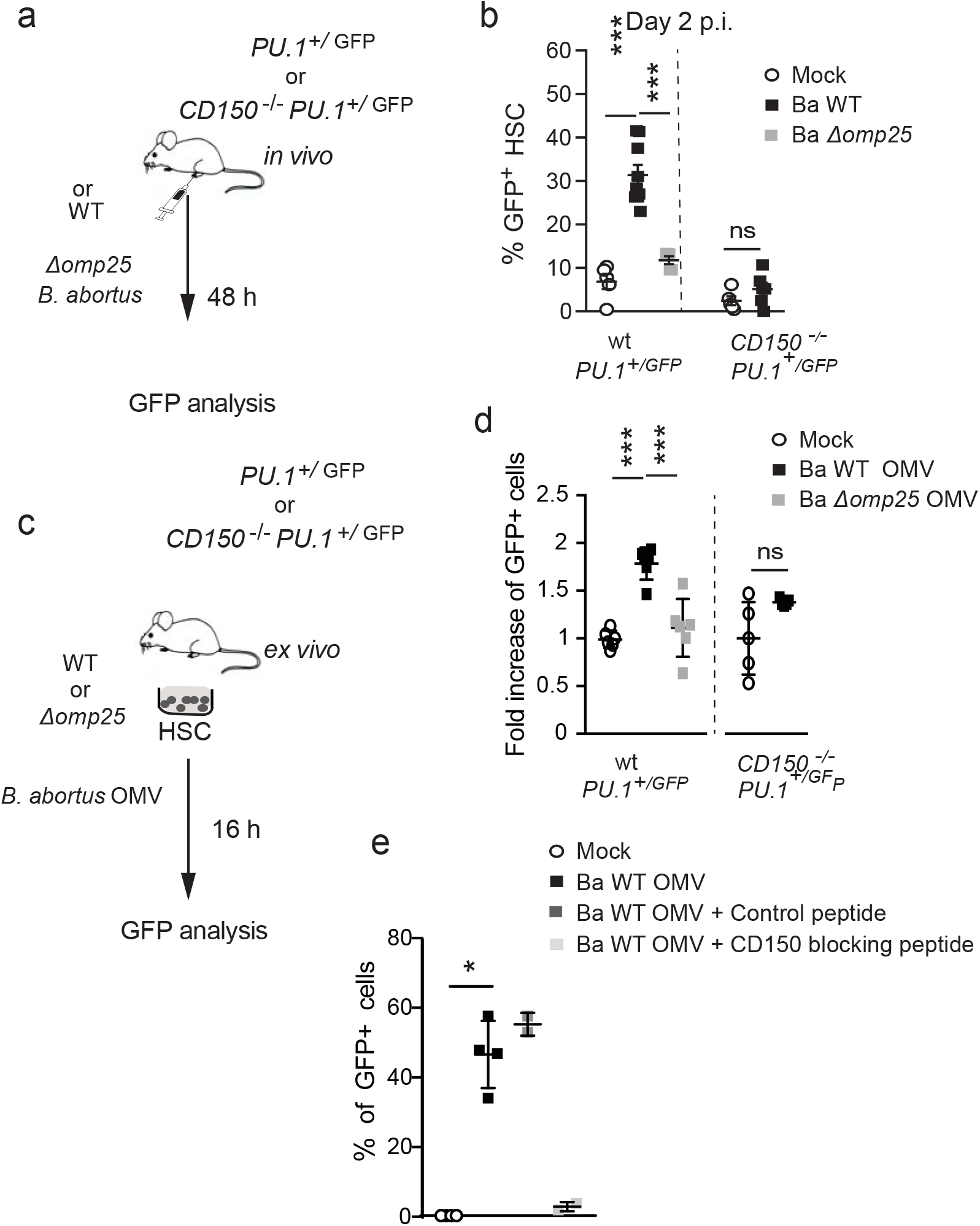
*Brucella* induces PU.1 upregulation in a Omp25/CD150 dependent manner. a) Experimental scheme *Pu.1*^+/*GFP*^ and *CD150*^-/-^ *Pu.1*^+/*GFP*^ mice were intraperitoneally injected with PBSx1 (Mock, ○) or inoculated with 1×10^6^ CFU of wild-type *B. abortus* (Ba WT, ■), Ba Δ*omp25* 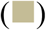. Two days after, the percentage of GFP expression in HSCs (lin^-^,Sca^+^,cKit^+^ CD48^-^, CD135^-^ CD34^-^) was assessed by Flow Cytometry. **b)** GFP expression in BM HSCs (lin^-^,Sca^+^,cKit^+^ CD48^-^, CD135^-^ CD34^-^) assessed by Flow Cytometry at D2 p.i.. Data (b) were obtained from distinct samples (from left to right, n=5; 10; 5; 5; 6) from 3 independent experiments. Mean ± SD is represented by horizontal bar. Significant differences from mock are shown. *** *P*< 0.001, ** *P*< 0.01, * *P*< 0.05. Absence of *P* value or ns, non-significant. Since data did not follow normal distribution, P-Value were generated using Kruskal-Wallis followed by Dunn’s test. **c)** Experimental scheme: HSC (lin^-^,Sca^+^,cKit^+^ CD48^-^, CD135^-^ CD34^-^) from *Pu.1*^+/*GFP*^ or *CD150*^-/-^ *Pu.1*^+/*GFP*^ mice were sorted, then stimulated *ex vivo* with PBSx1 or *B. abortus* WT OMVs, *B. abortus* Δ*omp25* OMVs. After 16h, the level of GFP in cells was assessed by Flow Cytometry. **d)** Fold change ratio of GFP expression in HSC 16 hours after *ex vivo* stimulation with *B. abortus* WT OMVs (**■**), *B. abortus* Δ*omp25* OMVs 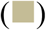 compared to mock (Mock, ○) assessed by Flow Cytometry (from left to right, n=6; 6; 6; 5; 5). **e)** Percentage of GFP^+^ HSC assessed by Flow Cytometry 0h (Mock, ○, n=4) or 16h after *ex vivo* stimulation with *B. abortus* WT (■, n=4), *B. abortus* WT OMVs and the control peptide (100μg/ml) (■, n=2) and blocking peptide of CD150 (100μg/ml) (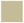, n=2). Data (e, f) were obtained from HSC of a pool of 3-4 mice, the pool of cells have been divided by the number of tested conditions. Each dot is a replicative experiment. Mean ± SD is represented by horizontal bar. Significant differences from mock are shown. ** P< 0.01, * P< 0.05. P-Value were generated using Mann Whitney test.

We then investigated the molecular mechanisms underlying the early response of HSC during *Brucella* infection. HSC can directly sense microbial compounds [14, 23, 24]. The CD150 receptor, known as one of the key markers of HSC [25], is also able to sense bacteria in dendritic cells and macrophages directly [12]. Indeed, in vitro studies showed that OmpC and OmpF of *E. coli* and *Salmonella spp*. binds to extracellular domain of CD150 [12]. Moreover, we have recently demonstrated that the outer-membrane protein 25 (Omp25) of *B. abortus* is a direct ligand of the extracellular domain of mouse CD150 in dendritic cells [13]. We asked whether HSC could detect *Brucella* via a direct Omp25/CD150 recognition. For this purpose we generated a new *Pu.1*^+/*GFP*^ reporter mouse model lacking CD150 (*CD150*^-/-^; *Pu.1*^+/*GFP*^). Infected *CD150*^-/-^ mice showed equivalent bacterial load in both spleen and BM during the onset and acute phase of infection (Supplementary Fig. 2). Moreover, infection did not affect either the total number of BM cells or the number of Lin^-^ cells in *CD150*^-/-^ mice (Supplementary Fig. 1). At 2 days post-infection, BM HSC from *CD150*^-/-^; *Pu.1*^+/*GFP*^ mice did not present any increase of GFP expression (Fig. 2b, right panel) in contrast to what we observed in *Pu.1*^+/*GFP*^ mice (Fig. 2b, left panel), suggesting that the induction of PU.1 by *B. abortus* is mediated by CD150.

To further test whether *B. abortus* directly binds CD150, PU.1 expression in HSC was analysed during infection with *B. abortus* WT and *B. abortus* lacking Omp25 (Ba Δ*omp25*) in *Pu.1*^+/*GFP*^ reporter mice (Fig. 2a). The upregulation of PU.1 observed in HSC in response to *B. abortus* WT infection was abolished when infected with Ba Δ*omp25* (Fig. 2b left panel), while bacterial CFU counts in spleen and BM were similar to those in mice infected with *B. abortus* WT (Supplementary Fig. 2). These data indicate that PU.1 upregulation in HSC during the onset of infection is dependent on Omp25/CD150 interaction.

To further demonstrate that the upregulation of PU.1 in HSC upon *Brucella* infection is due to the direct recognition of *B. abortus* Omp25 by CD150, we isolated HSC from the BM of *Pu.1*^+/*GFP*^ mice and treated them *ex vivo* with outer membrane vesicles (OMV) [26] from either *Brucella* WT or Ba *Δomp25* (Fig. 2c). At 16 h post-*ex vivo* stimulation of sorted HSC by *B. abortus* WT OMV, the number of GFP-expressing HSC increased two-fold (Fig. 2d), following the same trend as *in vivo* infection of *Pu.1*^+/*GFP*^ reporter mice (Fig. 2b). In contrast, upregulation of GFP-expressing HSC was abolished by either incubation of HSC with Ba *Δomp25* OMV (Fig. 2d) or with *B. abortus* WT OMV in the presence of CD150 blocking peptide (Fig. 2e) or in *CD150*^-/-^ (Fig. 2d). These data demonstrate that HSC can sense bacteria via a direct interaction of *Brucella* outer membrane protein Omp25 with CD150. This is the first demonstration of a direct recognition of a pathogen by HSC via CD150.

PU.1 upregulation in HSCs is a first sign of commitment towards the myeloid lineage [20, 27]. We therefore asked if the direct interaction between HSCs and *Brucella* induces a functional commitment of HSCs towards the myeloid lineage. We analysed the composition of HSC-downstream progenitors at 8 days post-infection in the BM. As expected, an increase in myeloid-biased MPP2/3 (Lin^-^, Sca^+^, cKit^+^, CD48^+^, CD135^-^) progenitors was observed in the BM of *wt* mice infected with *Ba* WT but not in the BM of *wt* mice infected with Ba *Δomp25* or infected mice lacking CD150 (Fig. 3a and Supplementary Fig. 1c for gating strategy). By contrast, the number of lymphoid-biased MPP4 (Lin^-^, Sca^+^, cKit^+^, CD48^+^, CD135^+^) was similar in infected and non-infected mice (Fig. 3b). Moreover, analysis of downstream committed progenitors revealed an increase of GMP and blood myeloid cells (Fig. 3c, d and Supplementary Fig. 1c for gating strategy). In addition, infection of *wt* mice by the Ba Δ*omp25c* complemented strain (Ba Δ*omp25* strain complemented with an Omp25-expressing plasmid) was able to rescue the increase of myeloid MPP2-3 and GMP demonstrating a direct role of Omp25 in controlling the increase of myeloid commitment via CD150 (Fig. 3a-c). Altogether, these data provide the first evidence that *B. abortus* induces an increase of myeloid cells production in a Omp25/CD150-dependent manner, though not at the expense of lymphoid cells.

**Figure 3:**
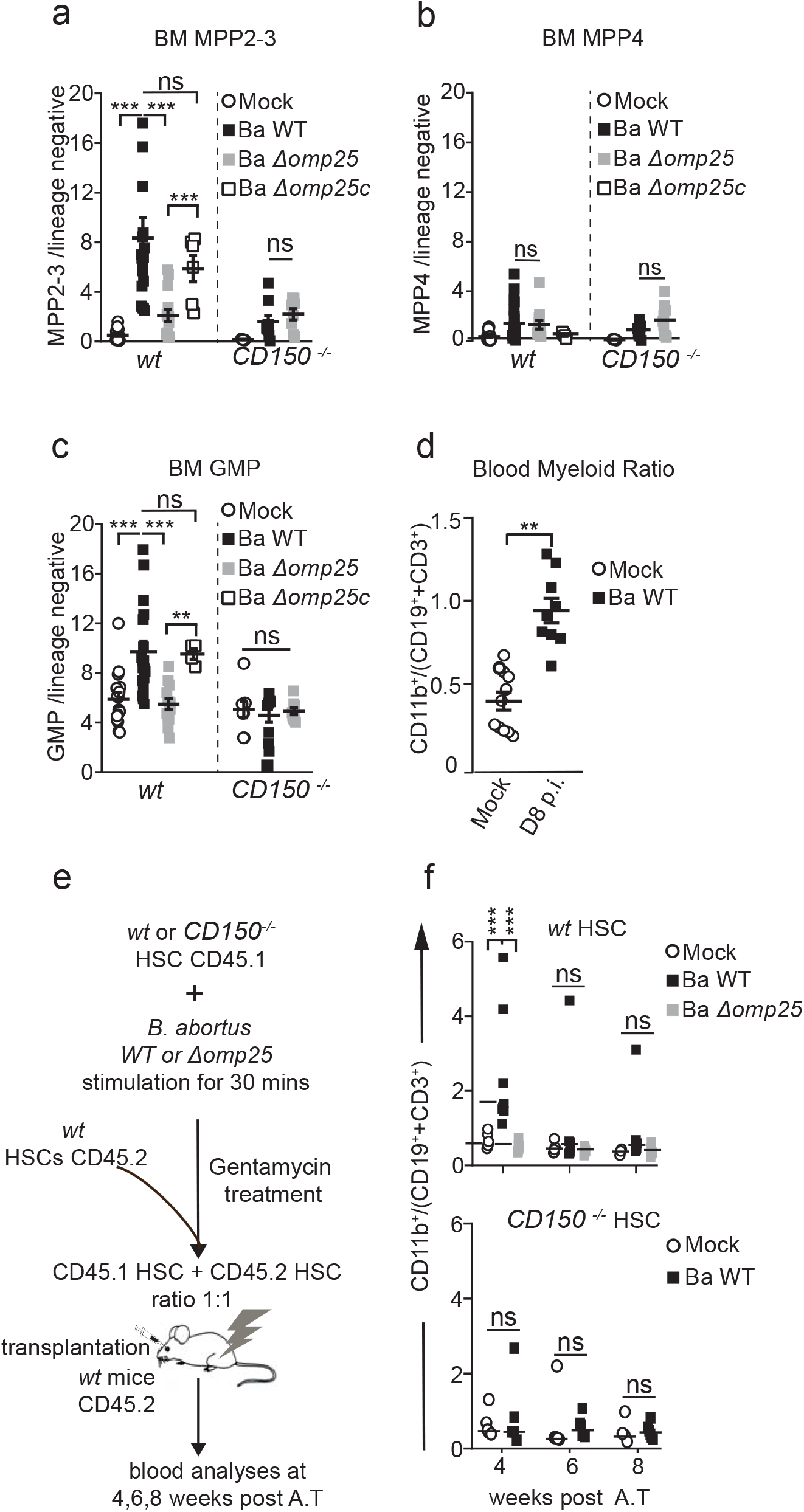
*B. abortus* induces HSC differentiation towards myeloid lineage. **a-d)** C57BL/6J wild-type (*wt*) and *CD150*^-/-^ mice were intraperitoneally inoculated with 1×10^6^ *B. abortus* CFUs. Eight days later, FACS analyses were performed for BM cells. Frequency of **(a)** MPP2-3 (lin^-^,Sca^+^,cKit^+^ CD48^+,^ CD135^-^) (from left to right: n=22; 19; 14; 6; 11; 9; 8), **(b)** MPP4 (lin^-^,Sca^+^,cKit^+^ CD48^+^ CD135^+^) (from left to right: n=22; 21; 12; 4; 9; 9; 9), **(c)** GMP (lin^-^,Sca^-^,cKit^+^ CD34^+^ CD16/32^+^) (from left to right: n=16; 18; 13; 4; 8; 9; 9), in Lin^-^ BM cells is shown for (Mock, ○) or inoculated with 1×10^6^ CFU of wild-type *B. abortus* (Ba WT, ■), *B. abortus* Δ*omp25* (Ba Δ*omp25* 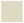) or *B. abortus* Δ*omp25* complemented with p:Omp25 (Ba *Δomp25c* □) mutants (the latter only for wt mice). **d)** Eight days post-infection, myeloid cells (CD45^+^, CD11b^+^) to lymphoid cells (CD3e^+^CD19^+^) ratio in blood is shown for (Mock, ○, n=11) or inoculated with 1×10^6^ CFU of *wild-type B. abortus* (Ba WT, ■, n=9), *B. abortus* Δ*omp25* (Ba Δ*omp25* 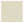). **e)** Experimental scheme: HSC from *wt* CD45.1 and *CD150*^-/-^ CD45.1 mice were sorted and then incubated *ex vivo* with *B. abortus WT* and *B. abortus* Δ*omp25* for 30 minutes. After 30 minutes, cells were washed and treated for 1 hour with gentamycin to kill extra-cellular bacteria. HSC were then transplanted into lethally irradiated *wt* CD45.2 recipients. FACS analyses of blood samples were performed at 4, 6 and 8 weeks post-transplantation. **f)** myeloid cells (CD45^+^, CD11b^+^) to lymphoid cells (CD3e^+^CD19^+^) ratio in CD45.1^+^ blood cells is shown for hematopoietic cells provided by CD45.1^+^ *wt* mice (upper panel) or *CD150*^-/-^ mice, (lower panel), non-infected (Mock, ○) or infected with 1×10^6^ CFU of wild-type *B. abortus* (Ba WT, ■), *B. abortus* Δ*omp25* (Ba Δ*omp25* 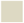) as described in e (from left to right, for WT n=12; 13; 10; 10; 9; 9; 12; 8; 8 and for CD150^-/-^ n= 9; 4; 14; 8; 11; 7). Data were obtained from distinct samples from 4 independent experiments (a-d) or from repetitive sampling from 2 (e-f) independent experiments. Mean ± SEM is represented by horizontal bar. Significant differences from mock are shown. *** P< 0.001, ** P< 0.01, * P< 0.05. Absence of P value or ns, non-significant. Since data did not follow normal distribution, P-Value were generated using Kruskal-Wallis followed by Dunn’s test.

HSCs are known to both self-renew and differentiate in order to replenish the whole hematopoietic system, properties that can be tested by transplantation in an irradiated host [6, 28]. In order to functionally confirm that the increased production of myeloid progenitors and mature cells was initiated by direct stimulation of HSCs by *Brucella*, we co-transplanted *ex vivo* stimulated and unstimulated HSCs in the same recipient. Towards this end we sorted HSCs (KSL, CD48^-^, CD34^-^, CD135^-^) from CD45.1 *CD150*^-/-^ or CD45.1 *wt* mice, stimulated them *ex vivo* with *Brucella* for 30 min and treated them with gentamicin to kill extracellular bacteria. We then transplanted the stimulated HSCs together with non-stimulated competitor HSCs (ratio 2:1) into lethally irradiated CD45.2 recipient mice (Fig. 3e). Blood analyses at 4 weeks post-transplantation show that HSCs stimulated with *B. abortus* WT generated more myeloid than lymphoid cells in the peripheral blood, compared to PBS-treated HSCs (Fig. 3f, upper panel and Supplementary Fig.3 for gating strategy). Again, the myeloid commitment bias of hematopoietic stem cells was abolished when HSCs were stimulated *ex vivo* with *B. abortus* Δ*omp25* (Fig. 3f, upper panel) or in *CD150*^-/-^ HSCs stimulated with *B. abortus* WT (Fig. 3f, lower panel). Furthermore, the increased myeloid to lymphoid ratio in the blood generated by *B. abortus* WT stimulated HSCs was transient and was not observed at 6- and 8-weeks post-transplantation. This indicates that HSCs re-equilibrate lineage commitment without compromising long-term multi-lineage contribution, as is also observed for direct M-CSF stimulation of HSC [20]. In summary, *in vivo* and *ex vivo* data prove for the first time that HSCs are able to sense directly live bacteria such as *Brucella abortus* via CD150 and transiently increase the production of myeloid cells in response.

Immune effector cell functions are globally altered in *CD150*^-/-^ mice [29]. We further examined whether, following infection with *Brucella*, the observed lack of HSC myeloid lineage commitment and downstream production of myeloid effectors in *CD150*^-/-^ mice is due to absent CD150-mediated bacterial recognition in HSCs or globally defective innate immune cell effector function. For this, we generated mixed hematopoietic chimera mice by transplanting a 1:1 mix of BM cells from *CD150*^+/+^; CD45.2 mice and *CD150*^-/-^; CD45.1 mice into irradiated CD45.2 recipient mice (Fig. 4a). This allowed investigation of both *CD150* HSCs and *CD150*^-/-^ HSCs in the same environment and in the presence of *wt* innate immune cells during infection. At twelve weeks post-transplantation, we infected chimeric mice with *B. abortus*. We then assessed their lineage output in both CD45.2 (*CD150^+/+^*) and CD45.1 (*CD150*^-/-^) compartments by analysing the myeloid/lymphoid ratio in mature blood cells and bone marrow progenitors (Fig. 4b-e). At 8 days post-infection with *B. abortus* WT, the blood myeloid/lymphoid ratio was higher in the *wt* compartment compared to the *CD150*^-/-^ compartment (Fig. 4b). The increase of blood myeloid/lymphoid ratio in the *wt* compartment was also abolished upon Ba Δ*omp25* infection of chimeric mice (Fig. 4b). In BM progenitor cells, the percentage of GMP (Fig. 4c) and myeloid-biased MPP2-3 cells (Fig. 4d) was also increased in the BM in a Omp25/CD150-dependent manner. Interestingly, Omp25/CD150 interaction did not perturb the number of lymphoid-biased multipotent progenitors MPP4 (Fig. 4e). These results confirm that myeloid commitment induced by *B. abortus* Omp25/CD150 interaction is intrinsic to HSC.

**Figure 4:**
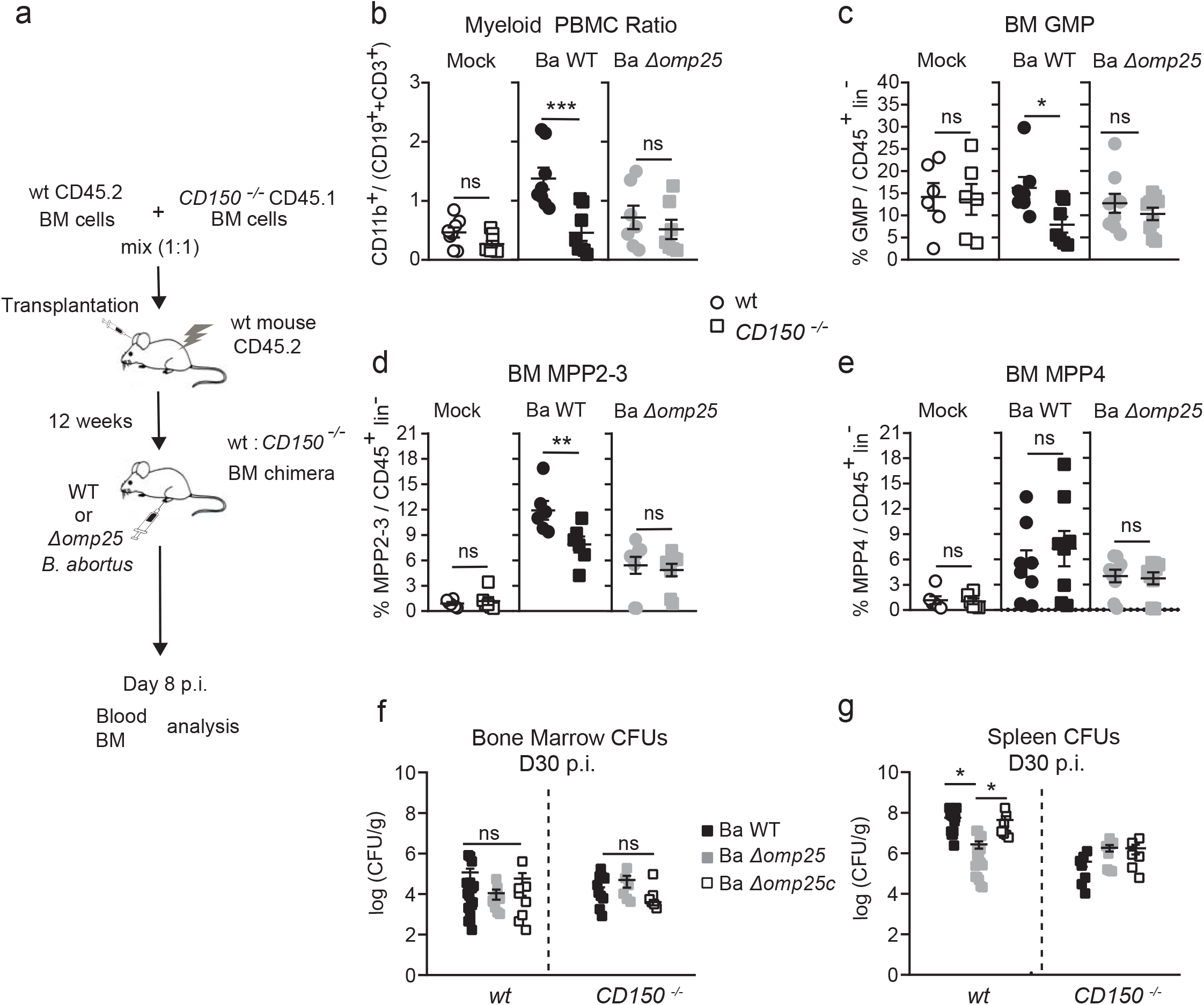
HSC myeloid bias induced by *Brucella* Omp25/CD150 interaction is hematopoietic cell autonomous. **a)** Experimental scheme: BM cells from *CD150*^-/-^ CD45.1 mice and *wt* CD45.2 mice were isolated from tibia and femur of mice and transplanted into lethally irradiated recipient mice. Twelve weeks after transplantation mice were intraperitoneally injected with PBSx1 (Mock, ○) or inoculated with 1×10^6^ CFU of wild-type *B. abortus* (Ba WT), *B. abortus* Δ*omp25* (Ba Δ*omp25*). Blood and BM were analysed 8 days after. **b)** Myeloid (CD45^+^,CD11b^+^ cells) to lymphoid (CD3e^+^ and CD19^+^ cells) in the blood (from left to right, n=8; 8; 8; 8; 7; 7); frequency of **c)** GMP (from left to right, n=6; 6; 7; 7; 9; 9), **d)** MPP2-3 (from left to right, n=6; 6; 6; 6; 9; 9) and **e)** MPP4 (from left to right, n=6; 6; 8; 8; 9; 9) in BM Lin-cells of wt (circle) and CD150^-/-^ (square) compartment is shown for chimeric mice intraperitoneal injected with PBS (Mock, non-filled symbols) or inoculated with 1×10^6^ CFU of wild-type *B. abortus* (symbol filled in black) *B. abortus* Δ*omp25* (symbol filled in grey) as described in a. **e-f)** CFU count per gram of organ at Day 30 post infection of WT and CD150^-/-^ mice for (e) BM (from left to right n=18; 9; 8; 12; 6; 6) and (f) spleen (from left to right n= 14; 18; 7; 7; 8; 6) infected with B. abortus WT (Ba WT ■), B. abortus Δomp25 (Ba Δomp25 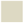) or B. abortus Δomp25 complemented with p:Omp25 (Ba Δomp25c □). Mean ± SEM is represented by horizontal bar. Significant differences from mock are shown. *** P< 0.001, ** P< 0.01, * P< 0.05. Absence of P value or ns, non-significant. Since data did not follow normal distribution, P-Value were generated using Kruskal-Wallis followed by Dunn’s test.

Infection by several pathogens is sometime associated with reduced red blood cell generation leading to a so called acute arrest of hematopoiesis (AAH) [30]. Interestingly, in the *wt* mice infected with *Brucella* the pourcentage of erythroid progenitors (MEP) in the BM decreased in a Omp25/CD150 dependant manner (Supplementary Fig.4a). To investigate if the reduced level of MEP results in anemia, we infected *wt* and *CD150*^-/-^ mice i.p. with *B. abortus*. At D8 p.i. we measured the hematocrit in the blood of the infected mice (Supplementary Fig.4b). As expected, the hematocrit of infected mice was reduced compared to non-infected mice in an Omp25/CD150-dependent manner (Supplementary Fig.4b).

These results indicate that in addition to increased myeloid commitment, reduced production of red cells already at level of progenitors cells is a Omp25/CD150-dependent feature of *Brucella* infection.

Notably, anemia associated with a decrease of red blood cells, hematocrit and haemoglobin was detected in 302 human patients (31.8% of men and 25% women) (Supplementary Fig.4c). These clinical data suggest that *Brucella* also alters red cell production in humans as in the mouse model leading to anemia and lack of body oxygenation. This phenomenea is responsible of fatigue, changes in metabolism and sometimes organ damage [31].

We also observed a splenomegaly in brucellosis patients, corroborating the results we obtained using the mouse model (Supplementary Fig 3d).

In addition, enhanced myeloid commitment has been shown to promote pathogen clearance of *E. coli* and *Salmonella* Typhimurium [17]. Nevertheless, for some pathogens, HSPC expansion is detrimental to the host and benefits the pathogen [32]. *CD150*^-/-^ mice are protected from *Trypanosoma cruzi* lethal challenge but are sensitive to *Leishmania major* and to attenuated *Salmonella* Typhimurium [31]. Bacterial load in the spleen of Ba Δomp25 *wt* infected mice or infected *CD150*^-/-^ mice at 4 weeks post-infection decreased compared to *wt* infected mice (Fig 4g). This suggests that the enhanced transient myeloid commitment induced by Omp25/CD150 benefits the bacterium. Indeed, *Brucella* infects and replicates in myeloid cells [7, 33]. This may be one of the strategies established by *Brucella* to promote chronic infection and help bacterial dissemination. The Omp25/CD150 axis can thus be considered as a new evasion strategy exploited by *B. abortus* to mediate its dissemination. Here, we present the novel finding demonstrating that CD150 is a bacterial sensor for HSC. How chronic activation of HSC by *B. abortus* could affect long-term function of HSC and the role of CD150 in HSC of patients experiencing a microbial challenge would be worth investigating in the future.

## METHODS

### Ethics

Animal experimentation was conducted in strict compliance with good animal practice as defined by the French Animal Welfare Bodies (Law 87–848 dated 19 October 1987 modified by Decree 2001-464 and Decree 2001-131 relative to European Convention, EEC Directive 86/609). INSERM guidelines have been followed regarding animal experimentation (authorization No. 02875 for mouse experimentation). All animal work was approved by the Direction Départementale Des Services Vétérinaires des Bouches du Rhône and the Regional Ethic Committee (authorization number 13.118). Authorisation of *Brucella* experimentation in BSL3 facility was given under the numbers: AMO-076712016-5, AMO-076712016-6 and AMO-076712016-7. All efforts were made to minimize suffering during animal handling and experimentation.

The study in humans was approved by the Ethics Committee of the Medical Faculty in Skopje, Republic of North Macedonia (No 03-7670/2).

#### Human study

The values of red blood cells, hematocrit and haemoglobin were retrospectively analyzed in 302 patients with human brucellosis before therapy was initiated. The patients were managed at the University clinic of infectious diseases and febrile conditions in Skopje from 2007 to 2018. Males were 217 and females 85 of them with a median age of 39 (range 3-79) years. The diagnosis of brucellosis was based on clinical findings compatible with brucellosis (arthralgia, fever, sweating, malaise, hepatomegaly, splenomegaly, signs of focal disease), confirmed by a qualitative positive Rose Bengal test and a Brucellacapt assay of >1/320. Hemoglobin thresholds used to define anemia were according to World Health Organization [World Health Organization (2008). Worldwide prevalence of anaemia 1993–2005 (PDF). Geneva: World Health Organization. ISBN 978-92-4-159665-7. Archived (PDF) from the original on 12 March 2009. Retrieved 2009-03-25].

### Mice

6-10 week-old female C57BL/6J mice from Charles River, *CD150*^-/-^ mice (kindly provided by Yusuke Yanagi) [29] or Pu.1^+/GFP^ mice [21, 22], both on a C57BL/6J background, were used. Animals were housed in cages with water and food *ad libitum* in the CIPHE animal house facility, Marseille. Two weeks before the start of experiments, mice were transferred to the BSL3, CIPHE, Marseille, and kept under strict biosafety containment conditions all along infection with live bacteria.

### Bacterial strains

*Brucella abortus* 2308 (Ba WT), *Brucella abortus* Δomp25 (kanR) (Ba *Δomp25*), or *Brucella abortus Δomp25c:pOmp25* (kanR, AmpR) (Ba *Δomp25c*) were used for infection. Ba Δ*omp25* was a gift from Pr. Ignacio Moriyón, University of Navarra.

### *Brucella* infection

Mice were inoculated intraperitoneally with 1×10^6^ CFU in 100μl of PBS for each *Brucella* strain. Strains were grown in Tryptic Soy Agar (Sigma Aldrich) for 5 days, then overnight at 37°C for 16 h under shaking in Tryptic Soy Broth (Sigma Aldrich) with kanamycin for Ba Δ*omp25* until the OD (OD at 600nm) reached 1.8 and with 25 μg/mL for the Ba Δ*omp25* strain or kanamycin and ampicillin 50 μg/mL for the Ba Δ*omp25pBBR4omp25* strain. All *Brucella* were kept, grown and used under strict biosafety containment conditions all along experiments in the BSL3 facility, Marseille. For Colony Forming Units (CFU) enumeration at different time points post-infection, spleen and bone marrow were collected [7]. Femur and tibia were flushed with 500 μl of ice-cold PBS to isolate BM cells. BM cell suspension was then plated in TSA plates. Spleens were collected and splenocytes were isolated by mechanical disruption.

Organs were harvested at 2, 8 or 30 days post-infection, weighted and then dissociated into sterile endotoxin free PBS. Serial dilutions in sterile 1xPBS were used to count CFU. Serial dilutions were plated in triplicates onto TSB agar to enumerate CFU after 3 days at 37°C.

### Transplantation assay

All donor cells were from *CD150*^+/+^ and *CD150*^-/-^ CD45.1 mice and transplanted into lethally irradiated (5,9 Gy) CD45.2 recipient mice. For competitive assays, mice were transplanted with equal numbers of 1×10^6^ total BM cells or 1×10^6^ lineage negative cells. For infected HSC transplantation, 1000 Sorted HSC (KSL, CD48^-^, CD135^-^, CD34^-^) were infected with Ba *WT* or Ba *Δomp25* at a MOI of 30:1. Bacteria were centrifuged onto cells at 400 g for 10 min at 15°C and then incubated for 45 min at 37°C under 5% CO2. Cells were washed twice with medium and then incubated for 1 h in medium containing 100 μg/ml gentamicin (Sigma Aldrich) to kill extracellular bacteria. Cells were than washed 3 times with PBS. Infected cells were mixed in a ratio (2:1) with non-infected HSC before transplantation. Haematopoietic reconstitution and lineage determination were monitored at 4 weeks, 6 weeks and 8 weeks post-transplantation in the peripheral blood. At 8 weeks post-transplantation mice were sacrified and tibia and femurs were harvested. BM cells were flushed from femur and tibia and resuspended in FACS media (PBS, 2%FCS, 5mm EDTA) for Flow Cytometry analyses.

### Flow Cytometry

For FACS sorting and analysis we used a FACSAriaIII or a LSR-X20 (BD) and the FlowJo software v10 (Treestar). For HSC and progenitor analysis, total BM cells were depleted of mature cells using a direct lineage depletion kit (Miltenyi Biotec) and stained with antibodies anti-CD34-APC or anti-BV421 (BD Bioscience, cloneRAM34), anti-CD135-PE-CF594 or anti-PE (BD Bioscience, clone A2F10.1), anti-CD150-PE-Cy7 or anti-BV711 (BioLegend, clone TC15-12F12.2), anti-CD117-BV605 (BioLegend, clone 2B8), anti-Sca-1-PrcpCy5.5 or anti-PE (ThermoFischer Scientist, clone D7), anti-CD48-BV510 or anti-PE-Cy7 (BD Bioscience, clone HM48-1), anti-CD16/32-PE or anti-APC-Cy7 (BD Bioscience, clone 2.4G2). When needed, anti-CD45.1-APC or anti-BV421 (BD Bioscience, clone A20) and anti-CD45.2-FITC or anti-PrcpCy5.5 (BD Bioscience, clone 104) were added. LIVE/DEAD (UV Fixable Blue Dead Cell Stain, ThermoFischer) was used as viability marker.

Blood cells were stained with anti-CD11b FITC (eBioescience, clone M1/70), anti-CD19-PE-Cy7 (BioLegend, clone 6D5), anti-CD45.2-PrcpCy5.5 (BD Bioscence, clone 104), anti-CD45.1-BV421 (BD Bioscience, clone A20), anti-CD3e-APC (BD Bioscience, clone 145-2C11) and anti-Ly6G-PE (BD Bioscience, clone 1A8). Red blood cells were lysed using BD FACS lysing solution (BD) for 10 min then fixed for 20 min with Antigen Fix, prior to acquisition.

### Haematopietic Stem Cells *ex vivo* challenge with *Brucella abortus* membrane extracts

All cultures were performed at 37°C under 5% CO_2_. Sorted HSC from wt or *CD150*^-/-^ mice were cultured in StemSpan SFEMII (Stem Cells) complemented with 50 ng/μL TPO (Peprotech) and 20 ng/μL SCF (Peprotech). Cells were stimulated with *Brucella* membrane extracts from Ba WT or with Ba *Δomp25* (10 μg/ml). *Brucella* membrane extracts were a gift from Pr. I. Moriyon, University of Navarra.

Sorted HSC were also cultured with blocking CD150 peptide (FCKQLKLYEQVSPPE, Auspep, 100 μg/ml) or control peptide (DLSKGSYPDHLEDGY, Auspep,100 μg/ml) (Thermo Scientific).

### Statistics

Results were evaluated by GraphPad Prism v8 software (GraphPad Software, San Diego, CA, USA) using. Statistical tests used are indicated in the figure legends. The value of *P < 0.05 was determined as significant.

## DATA AVAIBILITY

No datasets were generated during the current study.

## ACKNOWLEDGEMENTS

We thank all the staff of the CIML and CIPHE mouse houses, Atika Zouine, Marc Barad, and Sylvain Bigot of the CIML flow cytometry facility, Dr. Hervé Luche, Pierre Grenot of the CIPHE flow cytometry facility, Claude Napez and Philippe Hoest of the CIPHE BSL-3 facility. We acknowledge Pr. Ignacio Moriyon (University of Pamplona, France) for generous gift of mutant Brucella strains and Dr. Yusuke Yanagi and Dr.Masato Kubo for providing us with *CD150*^-/-^ CD45.1 mice. We thank Dr.Taymour Hamoudi from University of California San Francisco for critical reading of the manuscript and Dr. Sylvie Memet and Dr.Guillaume Hoeffel for discussions about this work.

This study was supported by institutional grants from the Institut National de la Santé et de la Recherche Médicale, Centre National de la Recherche Scientifique, and Aix-Marseille University to CIML and grants to JPG from Fondation pour la Recherche médicale (FRM grant number <u>DQ20170336745</u>), the Agence Nationale de Recherche/Investissements d’Avenir–Labex INFORM (ANR-11-LABX-0054) and Agence Nationale de Recherche/Investissements d’Avenir–A*MIDEX (ANR-11-IDEX-0001-02), to SS from ITMO Cancer Aviesan (Alliance Nationale pour les Sciences de la Vie et de la Santé, National Alliance for Life Science and Health) within the framework of the Cancer Plan (ISC19032ASA) and to M.H. Sieweke from the Agence Nationale pour la Recherche (ANR-17-CE15-0007-01 and ANR-18-CE12-0019-03), Fondation ARC pour la Recherche sur le Cancer (PGA1 RF20170205515), an INSERM-Helmholtz cooperation and the European Research Council (ERC) under the European Union’s Horizon 2020 research and innovation program (grant agreement number 695093 MacAge). LH received a fellowship from Investissements d’Avenir–Labex INFORM, BDL received a fellowship from the Fondation ARC. M.H. Sieweke was supported as a Berlin Institute of Health Einstein visiting fellow at MDC and is an Alexander von Humboldt Professor at TU Dresden.

## AUTHORS CONTRIBUTION

J.P.G, M.S, S.S conceived and J.P.G, S.S supervised the study. J.P.G, M.S, S.S, B.D.L, L.H, V.A design the experiments. L.H, G.G. and V.A performed all BSL-3 experiments and B.D.L and G.D performed experiments not requiring a BSL-3 facility. M.B was responsible of human studies. J.P.G, M.S, S.S, B.D.L, L.H, V.A interpreted the data. J.P.G, M.S, S.S, B.D.L, L.H, V.A wrote the manuscript

## DECLARATION OF INTERESTS

The authors declare no competing interests.

## FIGURES LEGENDS

**Supplementary 1:**
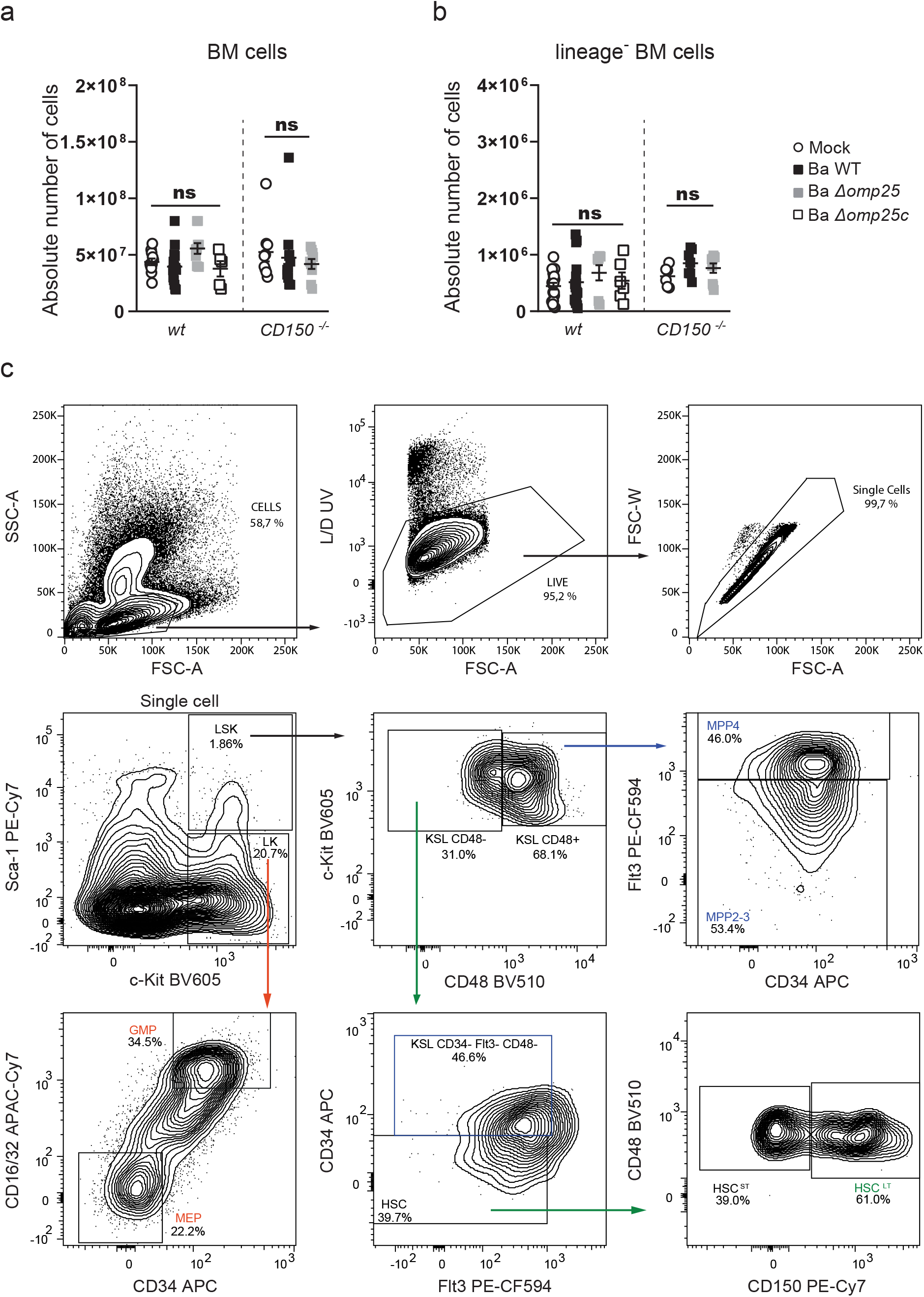
*Brucella* infection does not affect the number of lineage negative progenitors and BM cells. **a-b**) *wt* and *CD150*^-/-^ mice were intraperitoneally injected with PBS or inoculated with 1×10^6^ CFU of *B. abortus*. Eight days later, BM cells were isolated, cells were counted **(a)** and then depleted for mature hematopoietic cells as shown in Methods. Lin^-^ cells **(b)** were also counted for (Mock, ○) or infected *B. abortus* (Ba WT, ■), *B. abortus* Δ*omp25* (Ba Δ*omp25* 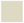) or *B. abortus* Δ*omp25* complemented with p:Omp25 (Ba *Δomp25c* □) mutants (the latter only for wt mice). From left to right, for BM, n=11; 14; 8; 5; 9; 11; 9 and for Lin-BM, n=18; 16; 8; 6; 6; 7; 9. Data obtained from distinct samples from 5 independent experiments. Mean ± SEM is represented by horizontal bar. Significant differences from mock are shown. *** P<0.001, ** P< 0.01, * P< 0.05. Absence of P value or ns, non-significant. Since data did not follow normal distribution, P-Value were generated using Kruskal-Wallis followed by Dunn’s test. c) Complete FACS gating for analysis of HSC and progenitors from lineage negative fraction of BM. First, cells are gated based on SSC/FSC and then single cells were selected. Viable cells were gated using UV Fixable Blue Dead stain. Described population are: LSK (lin^-^,Sca^+^,cKit^+^), HSCLT (lin^-^,Sca^+^,cKit^+^ CD48^-^, CD135^-^ CD34^-^,CD150^+^), MPP2-3 (lin^-^,Sca^+^,cKit^+^ CD48^+^, CD135^-^), MPP4 (lin^-^,Sca^+^,cKit^+^ CD48^+^, CD135^+^), GMP (lin^-^,Sca^-^,cKit^+^, CD34^+^, CD16/32^+^) and MEP (lin^-^,Sca^-^,cKit^+^, CD34^-^, CD16/32^-^).

**Supplementary 2:**
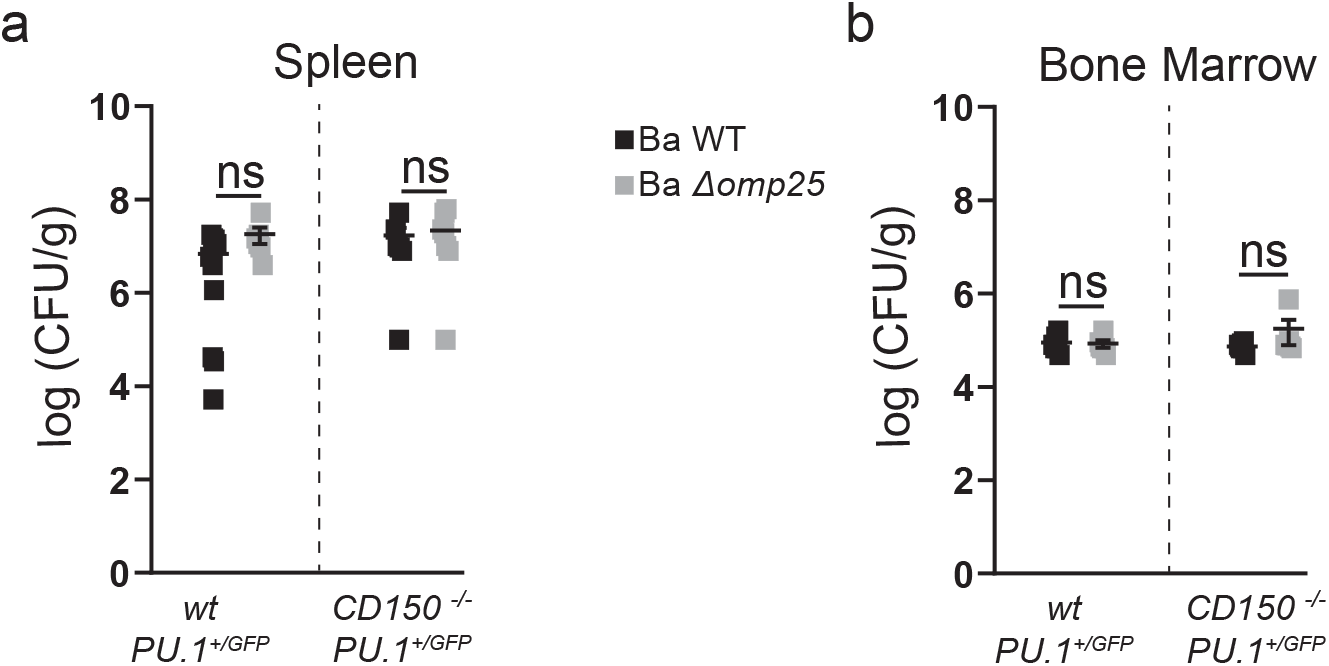
Infection burden in *Pu.1*^+/*GFP*^ mice. **a-b)** CFU count per gram of organ at Day 2 post infection as described in Fig 2.a. for **(a)** spleen (from left to right n=10; 6; 6; 8) and **(b)** BM (from left to right n=7; 6; 7; 7) of PBS injected (Mock, ○) or infected with *B. abortus* WT (Ba WT, ■), *B. abortus* Δ*omp25* (Ba Δ*omp25* 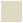) *Pu.1*^+/*GFP*^ and *CD150*^-/-^ *Pu.1*^+/*GFP*^ mice. Data obtained from distinct samples from 3 independent experiments. Mean ± SEM is represented by horizontal bar. Significant differences from mock are shown. *** P< 0.001, ** P< 0.01, * P< 0.05. Absence of P value or ns, non-significant. Since data did not follow normal distribution, P-Value were generated using Kruskal-Wallis followed by Dunn’s test.

**Supplementary 3:**
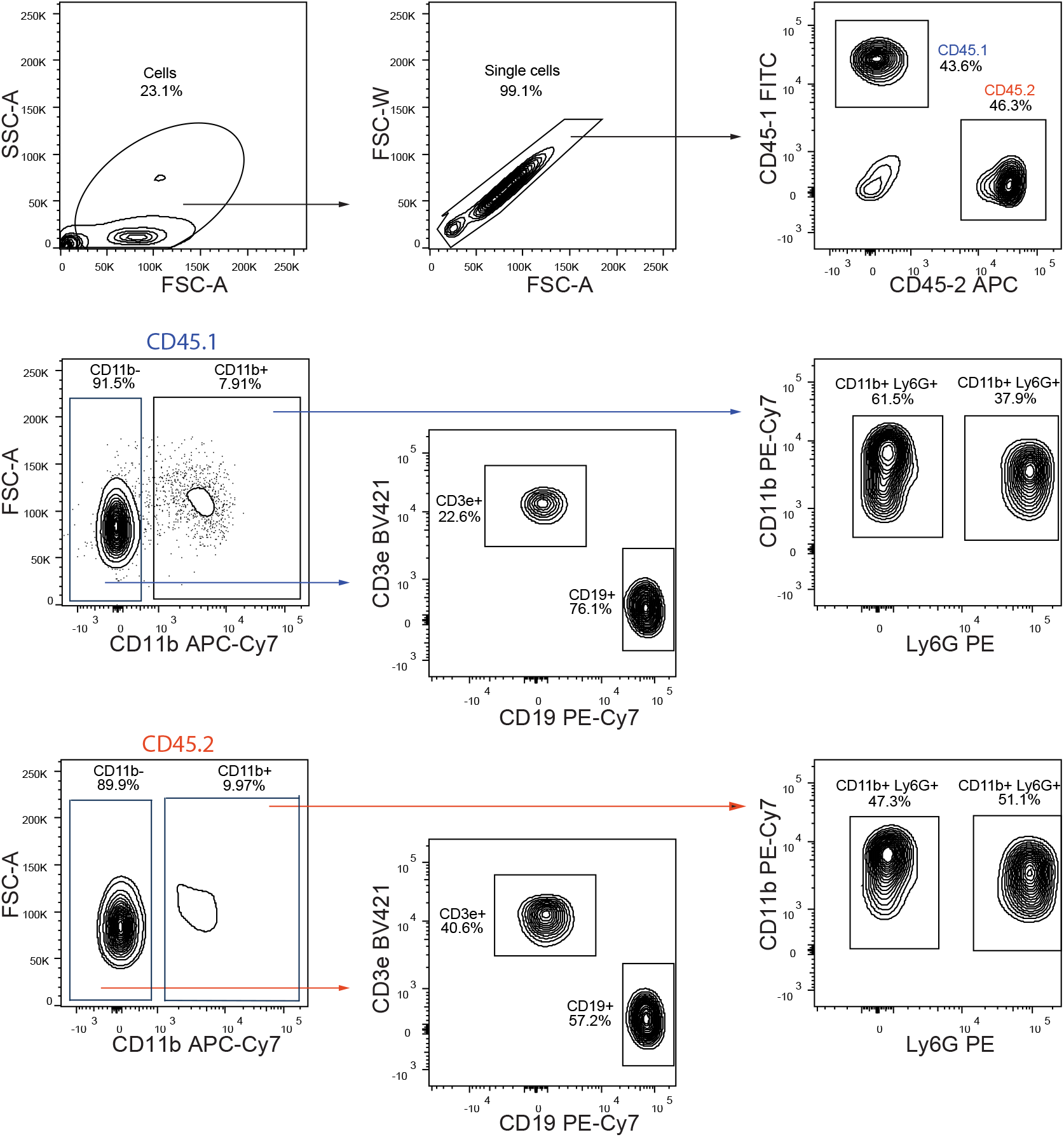
FACS gating for blood analysis from chimeric mice. Complete FACS gating for blood lineage output from chimeric mice. First, cells are gated based on SSC/FSC and then single cells were selected. CD150^-/-^ and WT cells were separated by gating on respectively CD45.1 and CD45.2 positive cells. In each, lymphoid cells were isolated by gating on CD11b negative cells and then, B cells are CD19 positive cells and T cells are CD3e positive cells. In Cd11b positive cells, granulocytes and monocytes are distinguished by gating onto respectively LY6G positive and negative cells.

**Supplementary 4:**
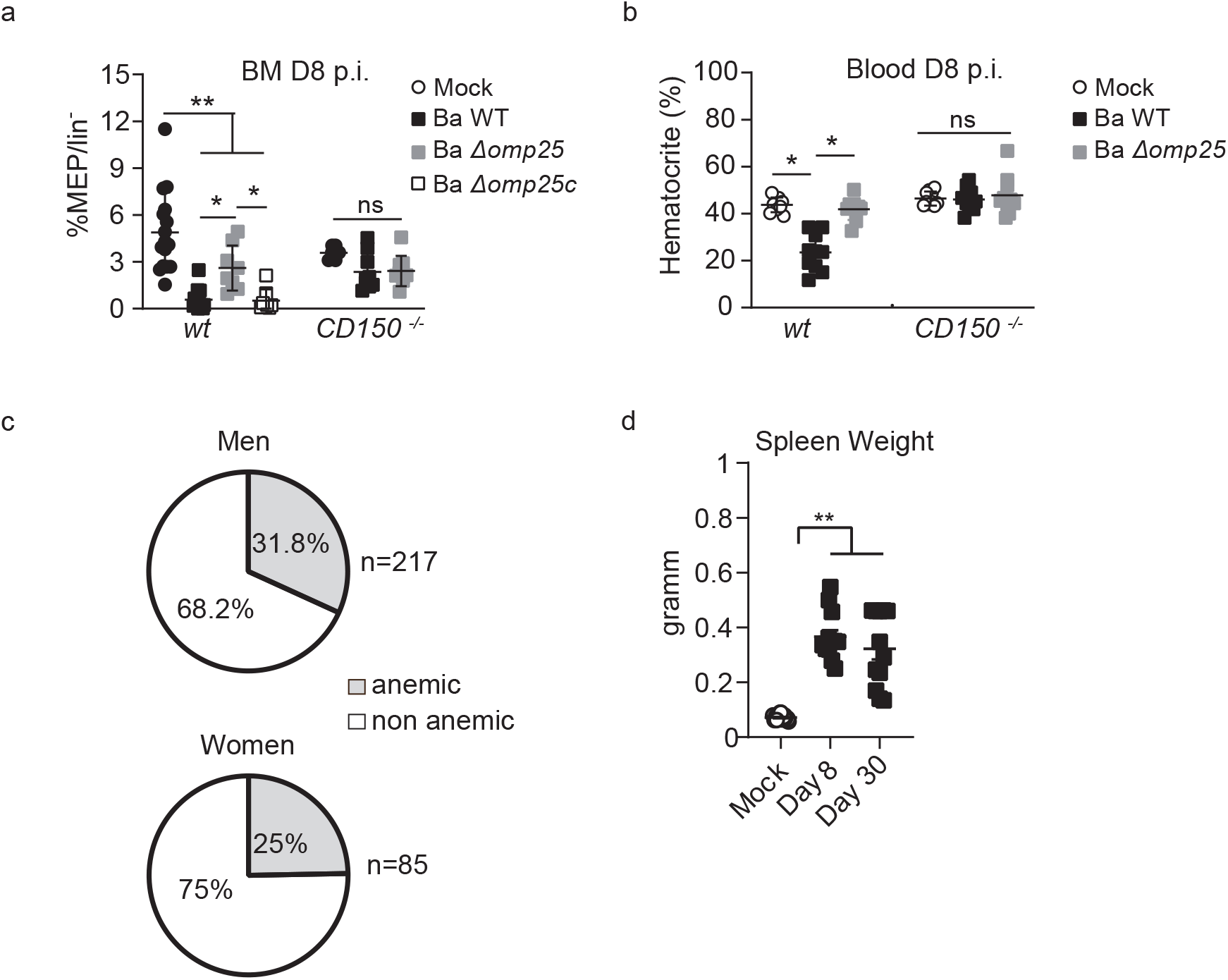
Brucellosis induces anemia. **a-b)** *<u>wt</u>* CD45.1 and *CD150*^-/-^ CD45.1 mice were intraperitoneally inoculated with 1×10^6^ CFU of *B. abortus*. BM cells were isolated from femur and tibia of the infected mice. a) Frequency of **MEP** (lin^-^,Sca^-^,cKit^+^ CD34^-,^ CD16/32^-^), in Lin^-^ BM cells is shown for (Mock, ○) or inoculated with 1×10^6^ CFU of wild-type *B. abortus* (Ba WT, ■), *B. abortus* Δ*omp25* (Ba Δ*omp25* 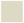) or *B. abortus* Δ*omp25* complemented with p:Omp25 (Ba *Δomp25c* □) mutants (the latter only for wt mice) (from left to right: n=16; 16; 8; 8; 8; 7; 9). **b)** At eight days post-infection, the percentage of haematocrit measured in blood is shown for (Mock, ○) or inoculated with 1×10^6^ CFU of *wild-type B. abortus* (Ba WT, ■), *B. abortus* Δ*omp25* (Ba Δ*omp25* 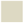) (from left to right, n= 7; 7; 6; 5; 8; 8). **c)** Percentage of brucellosis patients that present anemia before antibiotic treatment. Men upper panel and women lower panel. Anemia was characterized by a decrease of hematocrit, hemoglobin and erythrocytes (hematocrit <40% for men and <35% for women; hemoglobin <14g/dL for men and <12 g/dL for women, erythrocyte count <4 million for men and <3.8 million/mm3 for women). Data were obtained from distinct samples from 3 independent experiments (a-b), each with at least n=4 animals per condition, are shown and mean ± SEM is represented by horizontal bar. Significant differences from mock are shown. *** P< 0.001, ** P< 0.01, * P< 0.05. Absence of P value or ns, non-significant. Since data did not follow normal distribution, P-Value were generated using Kruskal-Wallis followed by Dunn’s test.

## Notes

### Competing Interest Statement

The authors have declared no competing interest.

## REFERENCES

1. Nebe, C.T., et al., Detection of Chlamydophila pneumoniae in the bone marrow of two patients with unexplained chronic anaemia. European journal of haematology, 2005. 74(1): p. 77–83.

2. Eldin, C., et al., 18F-FDG PET/CT as a central tool in the shift from chronic Q fever to Coxiella burnetii persistent focalized infection: A consecutive case series. Medicine, 2016. 95(34).

3. Allen, M.B., et al., First reported case of Ehrlichia ewingii involving human bone marrow. J Clin Microbiol, 2014. 52(11): p. 4102–4.

4. Hardy, J., P. Chu, and C.H. Contag, Foci of Listeria monocytogenes persist in the bone marrow. Dis Model Mech, 2009. 2(1-2): p. 39–46.

5. Reece, S.T., et al., Mycobacterium tuberculosis-Infected Hematopoietic Stem and Progenitor Cells Unable to Express Inducible Nitric Oxide Synthase Propagate Tuberculosis in Mice. J Infect Dis, 2018. 217(10): p. 1667–1671.

6. Weissman, I.L., Clonal origins of the hematopoietic system: the single most elegant experiment. The Journal of Immunology, 2014. 192(11): p. 4943–4944.

7. Gutierrez-Jimenez, C., et al., Persistence of Brucella abortus in the Bone Marrow of Infected Mice. J Immunol Res, 2018. 2018: p. 5370414.

8. Franco, M.P., et al., Human brucellosis. The Lancet Infectious Diseases, 2007. 7(12): p. 775–786.

9. Boettcher, S., et al., Endothelial cells translate pathogen signals into G-CSF-driven emergency granulopoiesis. Blood, 2014. 124(9): p. 1393–403.

10. Boettcher, S. and M.G. Manz, Regulation of Inflammation- and Infection-Driven Hematopoiesis. Trends Immunol, 2017. 38(5): p. 345–357.

11. Kiel, M.J., et al., SLAM family receptors distinguish hematopoietic stem and progenitor cells and reveal endothelial niches for stem cells. Cell, 2005. 121(7): p. 1109–21.

12. Berger, S.B., et al., SLAM is a microbial sensor that regulates bacterial phagosome functions in macrophages. Nat Immunol, 2010. 11(10): p. 920–7.

13. Degos, C., et al., Omp25 - dependent engagement of SLAMF1 by Brucella abortus in dendritic cells limits acute inflammation and favours bacterial persistence in vivo. Cellular Microbiology, 2020.

14. Takizawa, H., et al., Pathogen-Induced TLR4-TRIF Innate Immune Signaling in Hematopoietic Stem Cells Promotes Proliferation but Reduces Competitive Fitness. Cell Stem Cell, 2017. 21(2): p. 225–240 e5.

15. Mitroulis, I., et al., Modulation of Myelopoiesis Progenitors Is an Integral Component of Trained Immunity. Cell, 2018. 172(1-2): p. 147–161 e12.

16. Kobayashi, H., et al., Bacterial c-di-GMP affects hematopoietic stem/progenitors and their niches through STING. Cell Rep, 2015. 11(1): p. 71–84.

17. Takizawa, H., S. Boettcher, and M.G. Manz, Demand-adapted regulation of early hematopoiesis in infection and inflammation. Blood, 2012. 119(13): p. 2991–3002.

18. Pronk, C.J., et al., Tumor necrosis factor restricts hematopoietic stem cell activity in mice: involvement of two distinct receptors. Journal of Experimental Medicine, 2011. 208(8): p. 1563–1570.

19. Pietras, E.M., Inflammation: a key regulator of hematopoietic stem cell fate in health and disease. Blood, 2017. 130(15): p. 1693–1698.

20. Mossadegh-Keller, N., et al., M-CSF instructs myeloid lineage fate in single haematopoietic stem cells. Nature, 2013. 497(7448): p. 239–43.

21. Back, J., et al., PU. 1 determines the self-renewal capacity of erythroid progenitor cells. Blood, 2004. 103(10): p. 3615–3623.

22. Bryder, D., D.J. Rossi, and I.L. Weissman, Hematopoietic stem cells: the paradigmatic tissue-specific stem cell. The American journal of pathology, 2006. 169(2): p. 338–346.

23. Kobayashi, H., et al., Bacterial c-di-GMP affects hematopoietic stem/progenitors and their niches through STING. Cell reports, 2015. 11(1): p. 71–84.

24. Burberry, A., et al., Infection mobilizes hematopoietic stem cells through cooperative NOD-like receptor and Toll-like receptor signaling. Cell Host Microbe, 2014. 15(6): p. 779–91.

25. Kiel, M.J., et al., SLAM Family Receptors Distinguish Hematopoietic Stem and Progenitor Cells and Reveal Endothelial Niches for Stem Cells. Cell, 2005. 121(7): p. 1109–1121.

26. Boigegrain, R.A., et al., Release of periplasmic proteins of Brucella suis upon acidic shock involves the outer membrane protein Omp25. Infect Immun, 2004. 72(10): p. 5693–703.

27. Essers, M.A., et al., IFNalpha activates dormant haematopoietic stem cells in vivo. Nature, 2009. 458(7240): p. 904–8.

28. Till, J.E. and E.A. McCulloch, A direct measurement of the radiation sensitivity of normal mouse bone marrow cells. Radiation research, 1961. 14(2): p. 213–222.

29. Davidson, D., et al., Genetic evidence linking SAP, the X-linked lymphoproliferative gene product, to Src-related kinase FynT in T(H)2 cytokine regulation. Immunity, 2004. 21(5): p. 707–17.

30. Bi, L., et al., Acute arrest of hematopoiesis induced by infection with Staphylococcus epidermidis following total knee arthroplasty: A case report and literature review. Exp Ther Med, 2016. 11(3): p. 957–960.

31. Silverberg, D.S., et al., The pathological consequences of anaemia. Clin Lab Haematol, 2001. 23(1): p. 1–6.

32. Abidin, B.M., et al., Infection-adapted emergency hematopoiesis promotes visceral leishmaniasis. PLoS Pathog, 2017. 13(8): p. e1006422.

33. Salcedo, S.P., et al., BtpB, a novel Brucella TIR-containing effector protein with immune modulatory functions. Frontiers in Cellular and Infection Microbiology, 2013. 3.

